# Coalescence with background and balancing selection in systems with bi- and uniparental reproduction: contrasting partial asexuality and selfing

**DOI:** 10.1101/022996

**Authors:** Aneil F. Agrawal, Matthew Hartfield

## Abstract

Uniparental reproduction in diploids, via asexual reproduction or selfing, reduces the independence with which separate loci are transmitted across generations. This is expected to increase the extent to which a neutral marker is affected by selection elsewhere in the genome. Such effects have previously been quantified in coalescent models involving selfing. Here we examine the effects of background selection and balancing selection in diploids capable of both sexual and asexual reproduction (i.e., partial asexuality). We find that the effect of background selection on reducing coalescent time (and effective population size) can be orders of magnitude greater when rates of sex are low than when sex is common. This is because asexuality enhances the effects of background selection through both a recombination effect and a segregation effect. We show that there are several reasons that the strength of background selection differs between systems with partial asexuality and those with comparable levels of uniparental reproduction via selfing. Expectations for reductions in *N*_*e*_ via background selection have been verified using stochastic simulations. In contrast to background selection, balancing selection increases the coalescent time for a linked neutral site. With partial asexuality, the effect of balancing selection is somewhat dependent upon the mode of selection (e.g., heterozygote advantage vs. negative frequency dependent selection) in a manner that does not apply to selfing. This is because the frequency of heterozygotes, which are required for recombination onto alternative genetic backgrounds, is more dependent on the pattern of selection with partial asexuality than with selfing.

## Introduction

Coalescence theory forms the basis for much of the modern evolutionary inference made from genomic data. Although the coalescence of neutral alleles is simple in ideal populations, the process is altered by a myriad of biological realities such as population structure, changes in population size, and non-Poisson variance in reproductive success. Existing theory has tackled these and many other important deviations from the simplest case (WAKELEY 2008). In addition to ecological factors such as those above, selection at one site in the genome can profoundly influence the coalescence times of linked neutral sites, thereby affecting patterns of genomic diversity. Although selective sweeps can dramatically reduce coalescence times (MAYNARD SMITH and HAIGH 1974; KAPLAN *et al.* 1989), here we will focus on two other forms of selection: background and balancing selection.

Background selection refers to selection on deleterious alleles occurring in the genetic background of a focal neutral allele, and its resulting consequences for coalescence (reviewed in CHARLESWORTH 2012). Assuming selection is strong relative to drift, deleterious alleles are doomed to extinction and linked neutral sites will share the same fate unless they can escape via recombination (CHARLESWORTH *et al.* 1993). Looking backwards in time, this means that extant copies of a neutral site must trace their ancestry either avoiding deleterious backgrounds or recombining onto them only in the relatively brief period prior to their selective elimination. Thus, there are fewer potential ancestors for a neutral site than the actual number of individuals present in the preceding generations. In other words, background selection reduces the effective population size, shortening coalescent times. As this logic suggests, the consequences of background selection are much stronger if there is tight linkage between the focal neutral site and the locus experiencing deleterious mutations (HUDSON and KAPLAN 1995; NORDBORG *et al.* 1996a). Because of heterogeneity across the genome in the local deleterious mutation rate (e.g., due to variation in gene density and recombination rate), the strength of background selection is expected to vary across the genome. This is believed to be a major source of the variation in sequence diversity across chromosomes in species such as *Drosophila melanogaster* (CHARLESWORTH and CAMPOS 2014; ELYASHIV *et al.* 2014) and humans (MCVICKER *et al.* 2009).

Balancing selection preserves variation at a selected site, allowing two selected alleles to persist much longer than expected under drift alone. This means that coalescent times of the alternative selected alleles will be much longer than expected under neutrality. The same logic applies to two copies of a neutral site linked to alternative alleles at the selected site. Coalescence of more loosely linked neutral sites is less affected by the balancing selection that prevents coalescence at a selected site (HUDSON and KAPLAN 1988; KAPLAN *et al.* 1988; HEY 1991). By increasing coalescent times, balancing selection is expected to result in increased levels of neutral diversity at sites closely linked to targets of selection. Consistent with this, elevated neutral diversity is observed around loci strongly expected to be under balancing selection such as self-incompatibility alleles (RICHMAN *et al.* 1996; KAMAU and CHARLESWORTH 2005) and major histocompatibility loci (HUGHES and NEI 1988). In other cases, elevated levels of diversity have been used to identify potential candidates of balancing selection (ANDRÉS *et al.* 2009).

Most coalescent models implicitly assume that organisms are either haploid or diploid but obligately sexual and randomly mating. However, some of the most reproductively prolific diploids reproduce uniparentally much of the time via selfing or asexual reproduction (e.g., duckweeds, monogonant rotifers, ostracods, *Daphnia*, many grasses and fungi). Asexuality and selfing involve very different types of inheritance and, consequently, their “direct” effects on coalescent times are in opposite directions; asexuality increases coalescence time whereas selfing reduces it (see below). However, both forms of uniparental reproduction reduce the independence with which two alleles at separate loci within the same individual trace their ancestry backwards in time; that is uniparental reproduction increases the effective degree of linkage. As a consequence, uniparental inheritance is expected to magnify the importance of linked selection on coalescence.

Previous work has examined background and balancing selection for organisms that reproduce by both outcrossing and selfing (NORDBORG 1997). This work showed, for example, that the effect of background selection is stronger when selfing rates are high. Because effective population size can be dramatically reduced in selfers via background selection when considering genome-wide mutation, selfers suffer from concomitant reductions in the probability of fixation of beneficial mutations as well as increases in the probability of fixation of deleterious alleles (GLÉMIN and RONFORT 2012; KAMRAN-DISFANI and AGRAWAL 2014). This illustrates that studying the effect of linked selection on coalescent times of neutral sites is useful for understanding how uniparental reproduction affects adaptation via *N*_*e*_, in addition to its more obvious utility with respect to understanding levels of neutral diversity.

Here we develop coalescent models for organisms capable of both sexual and asexual reproduction. Like selfing, asexuality enhances the consequences of selection elsewhere across the genome, sometimes dramatically so. Partial asexuality and selfing are often lumped together, at least colloquially, because both are systems involving uniparental inheritance with reduced rates of genetic mixing. However, these alternative modes of uniparental reproduction heighten linked selection in somewhat different ways and we highlight the differences, though we do not present any original results with respect to selfing.

## Theory and Results

### Direct effects

In the absence of any linked selection, the expected coalescent time for two randomly sampled alleles in a diploid sexually outcrossing species is *E*[*T*] = 2*N* (WAKELEY 2008). For a species that produces a fraction *σ* of its offspring sexually and the remainder asexually, the expected coalescence time of two alleles sampled from different individuals is

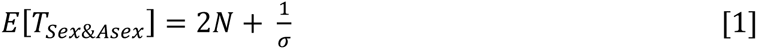

as shown by CEPLITIS 2003 (see also BENGTSSON (2003) and HARTFIELD *et al.* (2015) for further discussion of direct effects of asexuality).

As shown by NORDBORG and DONNELLY (1997), for a species that produces a fraction *o* of its offspring via outcrossing and the remainder by selfing, the expected coalescence time of two alleles sampled from different individuals is

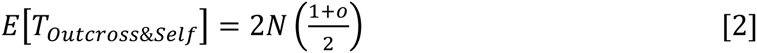

Contrasting [1] and [2], we see the direct effects of selfing and asexuality differs in several ways (Fig. 1). Increasing the rate of uniparental reproduction (i.e., reducing *o* or *σ*), reduces coalescent time in the case of selfing but has the opposite effect with partial asexuality. With selfing, coalescent time decreases linearly with declining rates of outcrossing. In contrast, the effect of partial asexuality on coalescent time is negligible unless the rate of sex is very small (i.e., the direct effect of partial asexuality can be ignored unless *σ* ∼ *O*(1/*N*)). Whereas very high selfing (*o* → 0) reduces the coalescent time by a maximum of 50%, the maximal increase in *E*[*T*] with very high asexuality (*σ* → 0) is unlimited in principle. However, if gene conversion is incorporated in the partial sex model, increases in coalescent time from asexuality are much more limited and can even be reversed (HARTFIELD *et al.* 2015).

**Figure 1.**
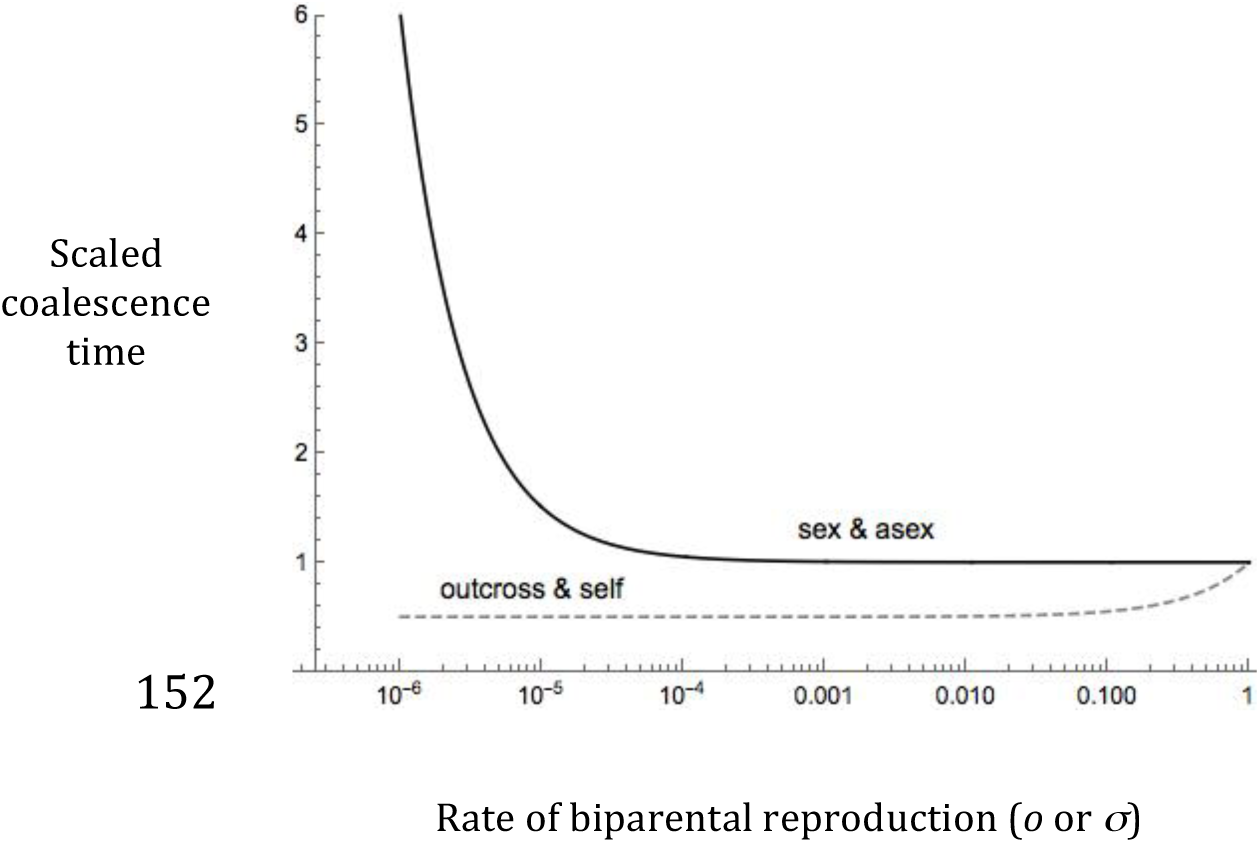
The direct effect of uniparental reproduction via asexuality or selfing on the scaled coalescence time (*E*[*T*]/2*N*). Partial asexuality has a negligible direct effect unless the rate of sex (*σ*) is on the order of the reciprocal of the population size; here *N* = 10^5^. High selfing reduces *N*_*e*_ to 50% of *N.*

We refer to the effects described above as the “direct” effects of uniparental reproduction to distinguish them from the “indirect” effects of uniparental reproduction in mediating the consequences of linked selection.

## Background Selection

The effects of background selection in sexual outcrossing species were first modeled in classic papers by HUDSON and KAPLAN (1994; 1995) and by NORDBORG *et al.* (1996a) using different approaches. To examine background selection in partially asexual species, we follow an approach similar to that used by NORDBORG (1997) who studied background and balancing selection in partial selfers. The main idea is that two sampled alleles can be found in a variety of different states (e.g., in the same or different individuals, in wild-type or mutant backgrounds). Based on transition probabilities, we can calculate the expected time to leave the current state and the probability with which the process will enter each of the other states, given that it has left its current state. If there are *k* states (other than coalescence), then we can create a system of *k* linear equations of the form

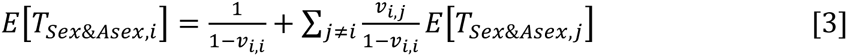

where *v*_*i,j*_ is the probability of the process transitioning from state *i* to state *j* (moving backwards in time). The system can be solved for the expected coalescent times from each state.

We make assumptions similar to those of previous models (HUDSON and KAPLAN 1994; 1995; NORDBORG *et al.* 1996a; NORDBORG 1997). We consider a single selected locus that can mutate from the wild-type *A*_1_ to the deleterious allele *A*_2_ at rate *μ*. The neutral locus is at a recombination distance *r* from the selected locus. Two neutral alleles could be in the same individual or two separate individuals. However, for each neutral allele we also need to consider its genetic background, both the selected allele present on the same chromosome as well as the selected allele on the homolog. We assume that *A*_1_/*A*_2_ individuals have a fitness of 1 – *hs* and that *A*_2_/*A*_2_ individuals have fitness 1 – *s* but are sufficiently rare that they can be ignored. The frequency of the deleterious allele is *q = μ/hs*. Because the allele frequency is assumed to be stable, this requires *N* is sufficiently large so that *Nq* >> 1. Technically, we assume 1/*N* ∼ *O*(*ξ*^3^), and *hs*, *q* ∼ *O*(*ξ*) where 0 < *ξ* << 1.

For fully sexual outcrossing species, diploid genotypes are transient so only haplotypes need to be considered. This “haplotype perspective” means that there are only three (non-coalesced) states in which two samples can exist: (i) both samples are found on *A*_1_-haplotypes, (ii) both samples are found on *A*_2_-haplotypes, and (iii) the two samples are on alternative haplotypes. With selfing there is the potential difficulty of needing to consider samples in the same or different individuals, potentially complicating the model with additional states. However, (NORDBORG 1997) used a separation of timescales argument to avoid this difficulty. With selfing, two samples from within the same individual either coalesce or transition into separate individuals at rates close to instantaneous with respect to coalescent times. Thus, it is possible to accommodate selfing into a system involving just three states (“the haplotype perspective”) by the appropriate adjustments of the transition probabilities.

In contrast, the separation of timescales assumptions is not applicable with partial asexuality as two samples may remain non-coalesced in the same individual for an extended period if rates of sex are low. Consequently, the system becomes considerably more complicated as we must keep an explicitly diploid perspective. Assuming homozygous mutant (*A*_2_*/A*_2_) genotypes are sufficiently rare that they can be ignored, there are eight (non-coalesced) states that must be considered: (i) both samples in a single *A*_1_*/ A*_1_ individual; (ii) both samples in a single *A*_1_*/A*_2_ individual; (iii) both samples in separate *A*_1_*/A*_1_ individuals; (iv) one sample in an *A*_1_*/A*_1_ individual and the other on the *A*_1_-haplotype of an *A*_1_*/A*_2_ individual; (v) one sample in an *A*_1_*/A*_1_ individual and the other on the *A*_2_-haplotype of an *A*_1_*/A*_2_ individual; (vi) both samples on the *A*_1_-haplotype of separate *A*_1_*/A*_2_ individuals; (vii) both samples on the *A*_2_-haplotype of separate *A*_1_*/A*2 individuals; and (viii) the samples in separate *A*_1_*/A*2 individuals, with one on an *A*_1_-haplotype and the other on an *A*_2_-haplotype. A complete model considering all the additional possibilities yields the same results [not shown]. The transition probabilities (*v*_*i,j*_) among these states, allowing for both sex and asexual reproduction, are given in the supplementary material.

If we assume that sex is common (*σ* ∼ *O*(1)) and linkage is tight (*r* ∼ *O*(*ξ*)), then the mean coalescence time is

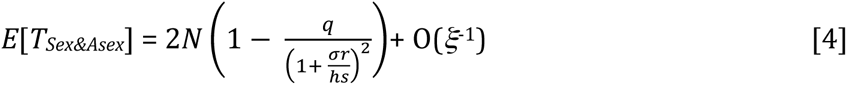

The term in parenthesis represents the reduction in the effective population size due to background selection. This result is equivalent to the classic result (Hudson & Kaplan 1994, 1995; Nordborg *et al* 1996a) but with the effective rate of recombination *σr* replacing *r* in the previous models, which assumed obligate sex. It is worth noting that partial asexuality (*σ* < 1) has an indirect effect on coalescence via background selection (represented by the parenthetical term in [4]), even though its direct effect of asexuality (as represented in [1]) is negligible in magnitude and not shown in this approximation. In this sense, the indirect effects of asexuality are much more important than the direct effects (unless rates of sex are very low, i.e., (*σ* ∼ *O*(1/*N*)). The result above is very similar to the result of (ROZE 2014) for partially asexual haploids. When the rate of sex is high, the shared history of homologs within a diploid are very brief relative to the coalescent time so it is unsurprising that the diploid model behaves like a haploid one.

If we assume that sex is rare but not too rare (*σ* ∼ *O*(*ξ*), *Nσ* >> 1) and that linkage is loose (*r* ∼ *O*(1)) then

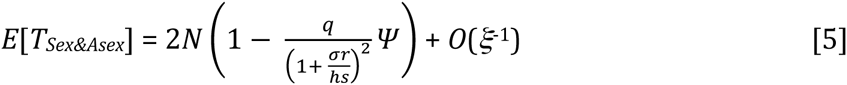

where

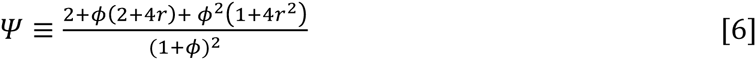

and *ϕ* ≡ σ/*hs* Under these assumptions, sex is still sufficiently common that the direct effect of asexual reproduction on coalescent time (via the effect highlighted in eq. (1)) is negligible. However, through the factor *Ψ* in [21], the reduced frequency of sex enhances the effect of background selection beyond the effect shown in [4], i.e., reducing *r* to *σr*. We now turn our attention to biologically interpreting *Ψ*.

This additional effect of low sex is clearer when we consider tight linkage (*r* ∼ *O*(*ξ*)) and even lower sex (*σ* ∼ *O*(*ξ*^2^)), then eq. [5] simplifies to

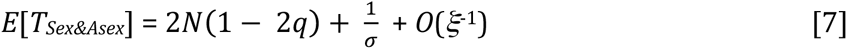

In this approximation, we also see the “direct” effect of sex on increasing the coalescent time, 1/*σ* (technically, [7] gives the coalescence time for two alleles sampled from separate individuals). The reduction in *N*_*e*_ due to background selection is captured by the factor (1 – 2*q*). This result is different than NORDBORG *et al.* (1996a) who, based on their result similar to eq. [4], showed that with no recombination the background selection term is (1 – *q*). The difference arises because the NORDBORG *et al.* (1996a) result assumes no recombination but obligate sex. Our result can be understood using the insight of CHARLESWORTH *et al.* (1993) that, without genetic mixing, the effective population size is reduced to mutation-free individuals. The frequency of mutation-free haplotypes is 1 – *q* but the frequency of mutant-free diploid genotypes is 1 – 2*q*. We can take [7] to mean that, in diploids with little sex, a focal neutral site is affected by the presence of a deleterious allele on its homolog just as much as it would be by one on its own chromosome, thus accounting for the 2*q* in place of *q*. With somewhat higher rates of sex, the neutral site is more likely to be freed by segregation from its homolog and thus will be less affected by mutations there.

The extent to which a focal site will be affected by mutations on the homolog with partial asexuality is captured by *Ψ*. This can be seen most easily when *r* is small (and *σ* ∼ *O*(*ξ*)) so that

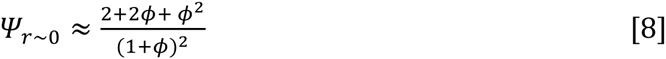

We can interpret *ϕ* as representing the odds that a neutral locus will escape a deleterious allele on the homolog before it is eliminated by selection because *ϕ* ≡ *σ* /*hs* represents the persistence time of a deleterious allele (1/*hs*) relative to the waiting time for sex (1/*σ*). From the equation above, it is easy to see that *Ψ* → 1 as *ϕ* → (i.e., *hs* << σ << 1) and *Ψ* → 2 as *ϕ* → ∞ (i.e., σ << *hs* << 1). In other words, background selection will depend as much on mutations on the homolog as on the focal chromosome (*Ψ* → 2) if the neutral site is unlikely to escape a deleterious allele on the homolog via sex before that allele is eliminated by selection.

Returning to the full version of *Ψ* in eq. [6], we find the somewhat surprising result that the sensitivity of background selection to mutation on the homolog *increases* with recombination, i.e., *∂Ψ*/*∂r* = 4 *ϕ* (1 + 2 r *ϕ*)/(1+*ϕ*)^2^ > 0. This can be understood as follows. When *r* = 0, the focal neutral site is guaranteed to escape the effects of a deleterious allele on the homolog via segregation if sex occurs. However, as *r* increases, the focal site is more likely to recombine onto the deleterious homolog during sex rather than escaping it. In fact, for unlinked loci (*r* = 1/2), we have *Ψ* = 2 (regardless of *ϕ*), indicating the focal site is just as strongly affected by mutations on the homolog as on its own chromosome.

In sum, asexual reproduction affects background selection in two ways: (i) via a “recombination effect” (reducing the effective rate of recombination from *r* to *σr*), and (ii) via a “segregation effect” (captured by *Ψ*, which mediates how much neutral alleles on one haplotype are affected by mutations on the homolog). Both of these effects on background selection occur even when the rate of sex is much higher than is required for a substantial direct effect of sex on coalescent time (i.e., *Nσ* >> 1). As a result of these effects, background selection can be much stronger in facultative asexuals; this is particularly noticeable at larger recombination distances (Fig. 2).

**Figure 2.**
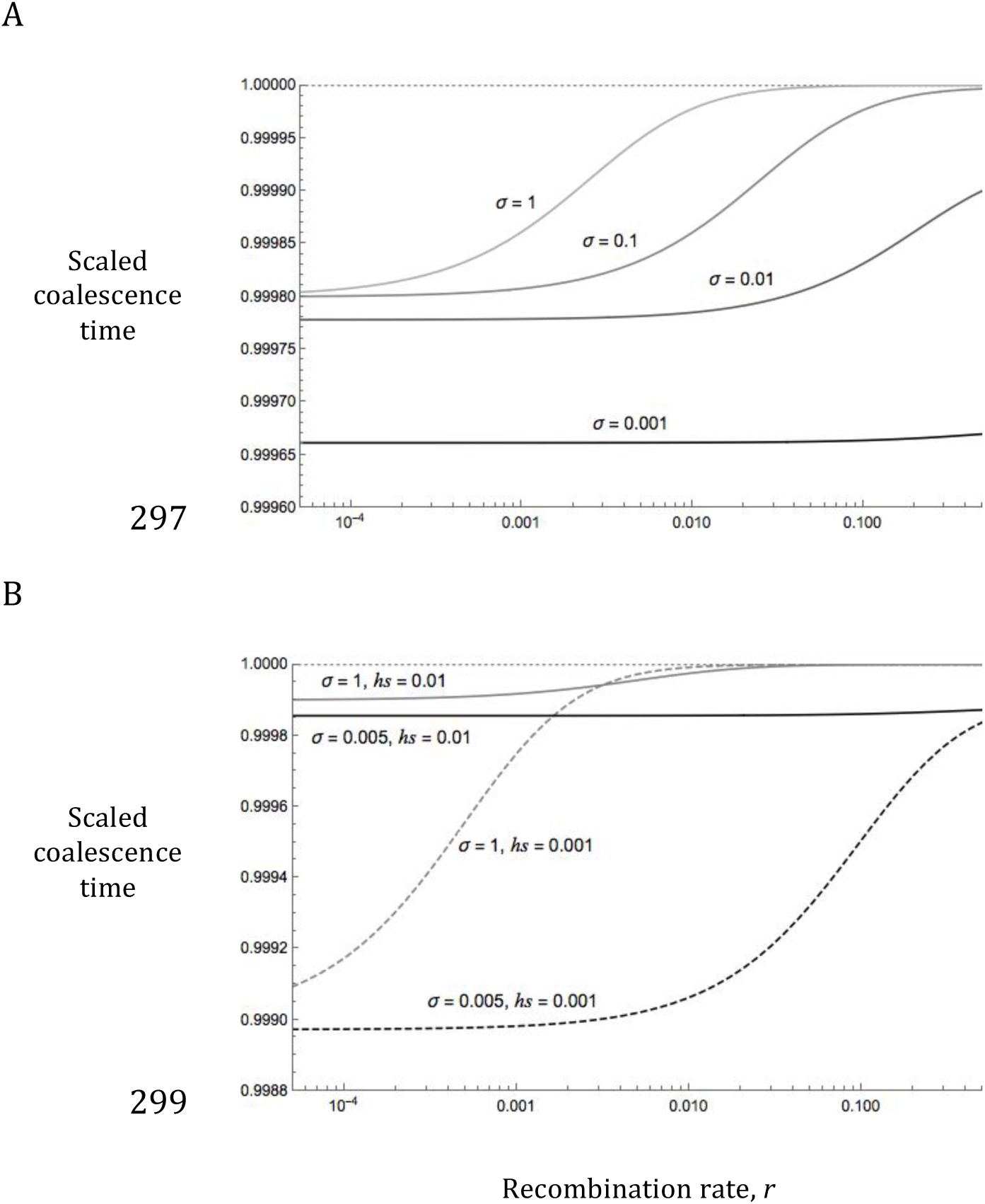
The scaled coalescence time (*E*[*T*]/2*N* or, equivalently, *N*_*e*_/*N*) as a function of the recombination distance between the focal neutral locus and the selected locus in the background selection model with partial asexuality. The reduction in coalescence times with low rates of sex is particularly at higher rates of recombination (A). The effects of background selection are larger with weak selection when recombination is low (B). Mutation rate: *μ* = 10^−6^.

### Extension to multiple loci

Following earlier models (HUDSON and KAPLAN 1995; NORDBORG *et al.* 1996a), we can extend the single-locus background selection results to many loci under the assumption of no disequilibria and no epistasis. Let *B*_*i*_ be the background selection factor for a single locus (i.e., the term in parenthesis of eq. [5]). Across many loci, the total effect of background selection is *B* = exp [∑ (1 – *B*_*i*_)]. Using eq. [5], we can calculate

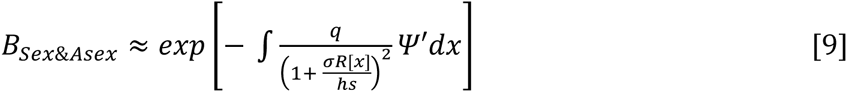

where the integration is over all positions *x* in the genome. *R*[*x*] is the recombination rate between the focal locus and the position *x* and *Ψ ′* is *Ψ* as given by [6] but replacing *r* with *R*[*x*].

Although [9] does not reduce to a simple analytical expression applicable to any genomic architecture, it can be evaluated numerically. Moreover, we can decompose the total background selection coefficient into components arising due to strongly linked, loosely linked, and unlinked genes: *B* = *B*^*TightlyLinked*^ *B*^*LooselyLinked*^ *B*^*Unlinked*^. Useful analytical approximations can be obtained for *B*^*TightlyLinked*^ and *B*^*Unlinked*^.

Here, we use “tightly linked” to refer to sites that are within a distance *m*_*max*_, measured in Morgans, of the focal site, where *m*_*max*_ is the maximum distance over which the map distance (*m*) is approximately linear with recombination rate. In other words, *m*_*max*_ is the maximum value of *m* where *m* ≈ *r*; for typical mapping functions, *m*_*max*_ < 0.05. We will consider a region 2*m*_*max*_ wide with the focal site at its center as “the tightly linked region” and assume deleterious mutations fall uniformly across this region. The background selection coefficient from the tightly linked region is

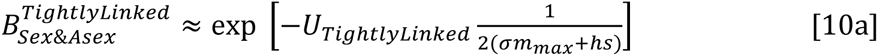

where *U*_*TightlyLinked*_ is the diploid mutation rate over the tightly linked region. For the case of *σ* = 1, this result is the same as eq. [8] of HUDSON and KAPLAN (1995). A much better approximation when *σ* < *hs* is

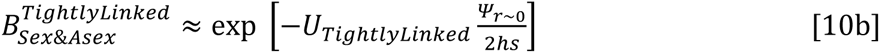

where *Ψ*_*r∼*0_ is given by [8].

Except in genomes with very small maps, most genes will be unlinked (*r* = ½) to the focal neutral site. Unlinked loci make a small contribution to the total effect of background selection in species where sex is high. With full sex (*σ* = 1) it has been shown by other means (CHARLESWORTH 2012) that

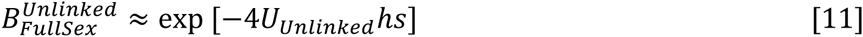

where *U*_*Unlinked*_ is the diploid rate of deleterious mutation over all sites unlinked to the neutral focal site. However, when sex is low (*σ* << 1), then the background selection coefficient due to unlinked sites based on [5] and [9] is

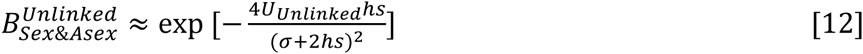

where *U*_*Unlinked*_ is the diploid rate of deleterious mutation over all sites unlinked to the neutral focal site. It is clear from eqs. [10-12], that the effects of background selection are much greater when sex is low, especially the contribution from unlinked loci.

To explore background selection further, we will assume that the focal chromosome exists in the center of a 1 Morgan chromosome and that the total map length of the genome is *L* Morgans (where *L* ≥ 1). Deleterious mutations are uniformly distributed across the genome, i.e., *μ* = ½ *U*/*L* where *U* is the diploid genome-wide mutation rate. This means that *U*_*TightlyLinked*_ = 2*Um*_*max*_/*L* and *U*_*Unlinked*_ = *U*(*L* – 1)/*L*. The background selection coefficient due to loosely linked (*m*_*max*_ < *r* < ½) sites is

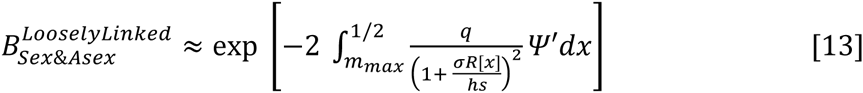

This integral can be expressed in closed form using Haldane’s mapping function (*R*[x] = exp[1 – 2*x*]/2) but the expression is rather complicated and is left for the supplementary material.

Figure 3 illustrates that the genome-wide effects of background selection can be substantial when *σ* < 1, even in genomes with long maps (such that gene density is low). If sex is low (*σ* < 1%), effective population size may be reduced by one or more orders of magnitude with relatively mild mutation rates (*U* = 0.1). In a moderate to large population (*N* > 10^4^), there would be a negligible direct effect on coalesce time of asexual reproduction with *σ* = 10% or 1%, (Bengtsson 2003; Ceplitis 2003; Hartfield et al 2015), but the indirect effect through background selection can be large. The effect of background selection is more sensitive to the strength of selection in long maps than in short maps, with stronger selection resulting in larger effects of background selection, provided *σ* is not too low. With lower sex, a larger proportion of total background selection is attributable to unlinked loci (Fig. 3B). This proportion is greater in larger genomes (as more loci will be unlinked) and when selection is strong, as strongly selected loci do not need to be closely linked to exert their effects (NORDBORG *et al.* 1996a).

**Figure 3.**
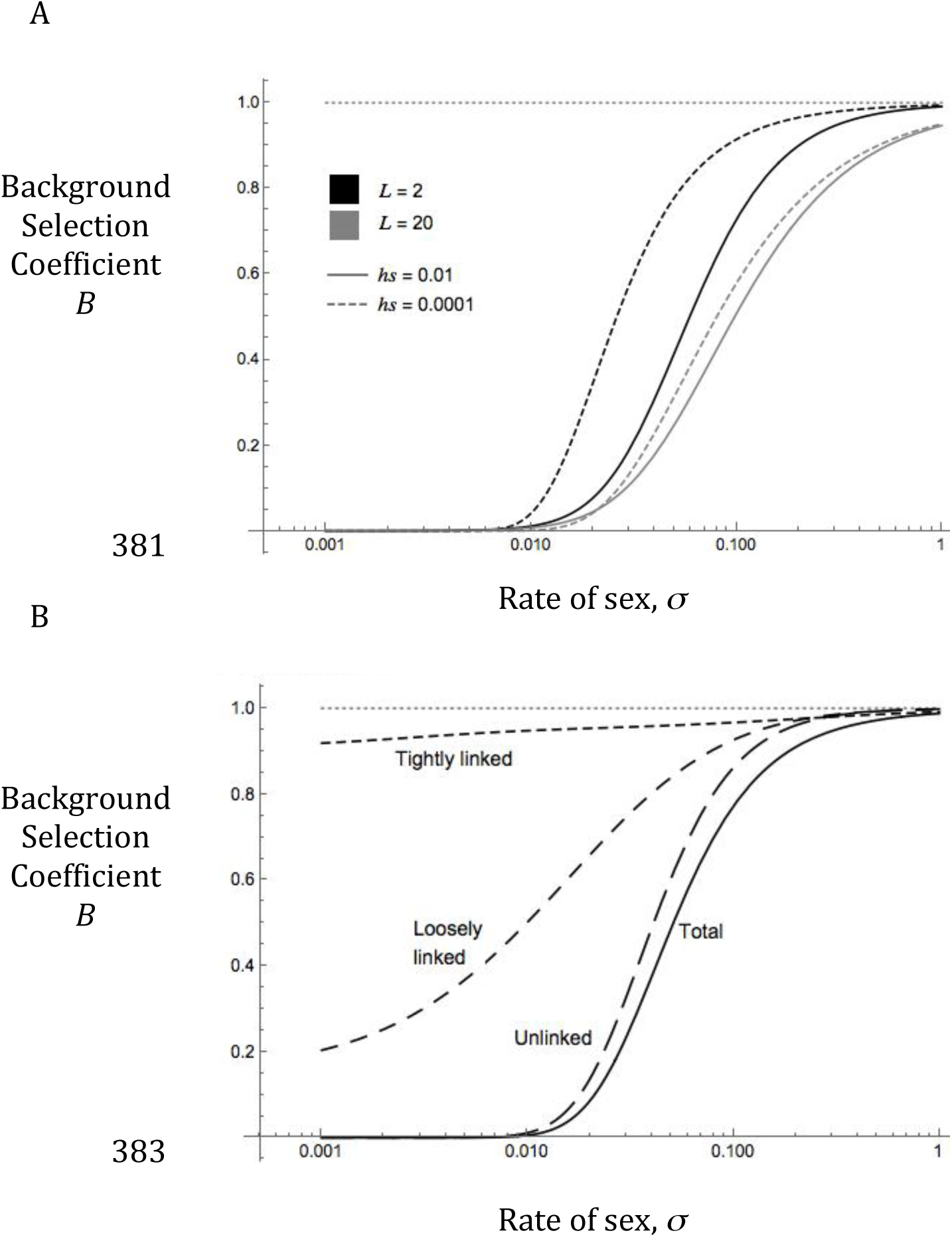
Genome-wide background selection in systems with sexual and asexual reproduction. (A) The background selection coefficient *B* as a function of the rate of sex considering mutations across the entire genome. Results for genomes with relative short and long maps (*L* = 2 vs. 20 Morgans) are shown. The longer map implies lower gene density as mutation rate is held constant (*U* = 0.1); mutations are assumed to be uniformly distributed across the genome. (B) The background selection coefficient for different genomic regions (tightly linked, loosely linked, and unlinked sites) as well as the total; parameters *L* = 10, *hs* = 0.005, *U* =0.1. Tightly linked loci make the largest contribution to background selection when rates of sex are very high but unlinked loci make the largest contribution when sex is rare.

### Simulation Results

The extrapolation to multiple loci used in [9] and in previous work (HUDSON and KAPLAN 1995; NORDBORG *et al.* 1996a; GLÉMIN and RONFORT 2012) relies on the assumption that loci are independent (i.e., no linkage disequilibrium) and that each selected locus is close to its deterministic equilibrium (*q* = *μ/hs*). These assumptions become increasingly suspect as the effective rate of recombination declines and as background selection becomes sufficiently strong that *N*_*e*_ (≈ *BN*) is substantially reduced. When *BNhs* is predicted to be low (< 1), then we expect that analytical approximations will overestimate the strength of background selection. Under these conditions the analytical approximation is expected to overestimate the reduction in *N*_*e*_ due to background selection, presumably because when drift is strong then polymorphism is lost from selected sites, i.e., *q* = 0 < *μ/sh*, making background selection weaker than predicted. Previous simulation studies have observed that analytical approximations tend to overestimate the strength of background selection when selection is weak or recombination is low (NORDBORG *et al.* 1996a; KAISER and CHARLESWORTH 2009; KAMRAN-DISFANI and AGRAWAL 2014).

To test the analytical approximations presented here, we extended the simulation of (KAMRAN-DISFANI and AGRAWAL 2014) to consider a population where individuals can reproduce asexually with a probability *σ*, via Wright-Fisher dynamics. We measured *N*_*e*_ by tracking a linked quantitative trait over time; when an individual reproduced, the value of this trait was perturbed by a value drawn from a normal distribution with mean 0 and variance 1. The steady-state variance of this trait is equal to the effective population size (LYNCH and HILL 1986; KEIGHTLEY and OTTO 2006).

For a given net mutation rate *U* and heterozygote selection coefficient *sh*, mutations were initially distributed amongst the population according to mutation-selection balance. Unless stated otherwise, the population then reproduced for *5N* generations, to ensure that neutral markers reached a steady state (KEIGHTLEY and OTTO 2006). The population-wide *N*_*e*_ was then measured at 200 equally spaced intervals for a further *5N* generations. The simulation was repeated for 192 burn-ins. The mean *N*_*e*_ from each time series was determined, and simulation points are calculated as the means of these values along with 95% confidence intervals. In general there is a good match between simulation results and analytical approximations (Fig. 4A). The effects of background selection tend to be overestimated if sex is low (*σ* ∼ 0.001), where background selection is predicted to be very strong.

**Figure 4:**
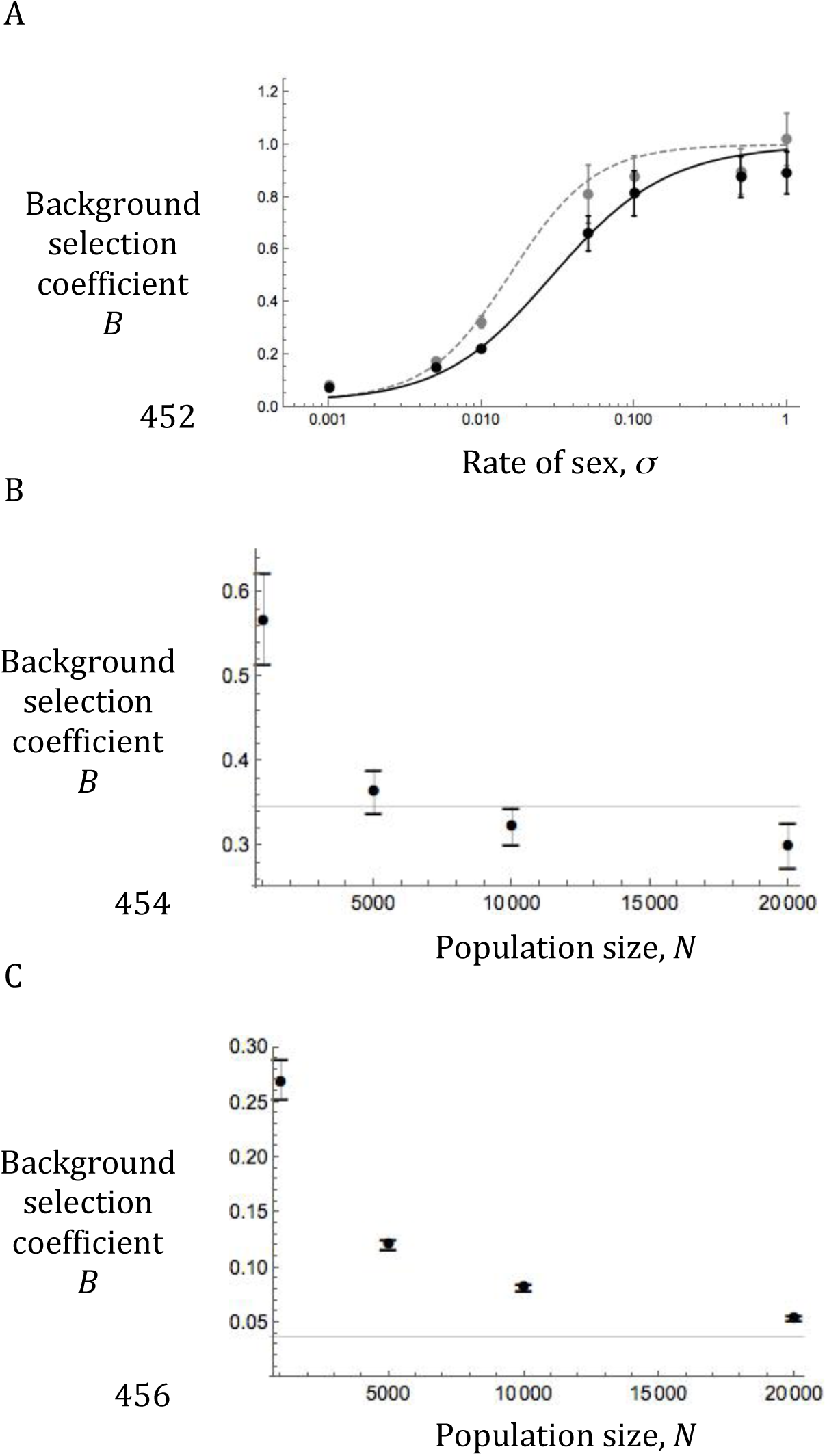
Simulation comparisons of background selection. (A) Simulation results (points) and analytical approximations (lines) of *B* = *N*_*e*_ /*N* as a function of the rate of sex. Parameters are *U* = 0.02, *s* = 0.02, *h* = 0.25, and *L* = 1 (black) or *L* = 10 (grey). Bars on simulation points represent 95% confidence intervals. (B) Background selection effect for *σ* = 0.01 as a function of the population size. Parameters are *U* = 0.02, *s* = 0.02, *h* = 0.25, and *M* = 10. The horizontal black line is the theoretical expectation given with Equation 9. (C) Same as (B) but for *σ* = 0.001.

Because the analytical approximation for *B* is expected to fail as *BNhs* becomes small, we examined the correspondence between simulations and the predicted value of *B* for different population sizes. As expected, the observed value of *B* is much larger than the analytical approximation (i.e., the strength of background selection is overestimated by [9]) when *N* is small but the discrepancy declines with larger *N* (Fig. 4B, C). This implies that the analytical approximations work well provided *N* is ‘sufficiently’ large. Though imperfect, the approximations offer good insight into the effects of background selection. Nonetheless, it is important to remember that the extremely strong effects of background selection predicted under low rates of sex (e.g., *B* < 10^−5^) are unlikely to be realized unless applied to species with a massive census size. Though the most obvious problem is that the analytical approximation overestimates background selection when *N* is not sufficiently large, we also find instances where the analytical approximation underestimates the strength of background selection when *N* is large (Fig. 4B). This could suggest a weak effect arising from multi-locus associations not captured in the extrapolation of the single-locus analysis to genome-wide background selection given by [9]. This effect does not appear to be sensitive to *h* (not shown) and so is unlikely to be a direct result of segregation load.

### Comparison with selfing

Another mode of uniparental reproduction, selfing, also enhances the effect of background selection. As shown by NORDBORG (1997), the coalescent time with selfing is

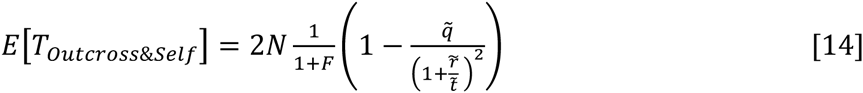

Here *F* is the inbreeding coefficient which can be written as function of the rate of outcrossing *F* = (1 – *o*)/(1 + *o*). The fraction 1/(1+*F*) is the direct effect of selfing on reducing *N*_*e*_ (shown in [2] as (1 + *o*)/2), and will not be considered further. The term in parenthesis in [14] is the background selection coefficient. With pure outcrossing the background selection coefficient is the same as the parenthetical term from [4] with *σ* = 1. Relative to this, selfing changes background selection in two ways. First, it increases the average strength of selection against a deleterious mutation from *sh* to 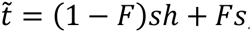, where *s* is selection against the mutation in the homozygous state. This also changes the equilibrium frequency of the deleterious allele to 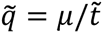. More importantly, selfing reduces the effective rate of recombination from *r* to 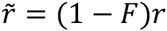. When selfing rates are high (*o* << 1), the background selection coefficient simplifies to

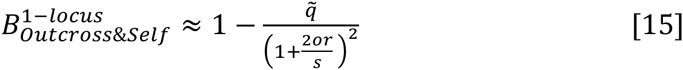

There are three important differences between this result with the equivalent term for background selection in partial asexuals [5]. First, the segregation effect captured by *Ψ* effectively doubles the relevant mutation rate for partial asexuals relative to selfers. Second, a bit of outcrossing in a highly selfing system results in effectively twice as much recombination as the equivalent amount of sex in a highly asexual system (i.e., *σr* in [5] vs. 2*or* in [15]). In sex/asex systems, recombination can only occur with sexual reproduction so the effective recombination rate depends directly on *σ*. In outcross/self systems, recombination can happen during the production of selfed or outcrossed progeny. However, the only parental genotypes in which recombination is relevant are double heterozygotes, which are created at a rate proportional to the rate of outcrossing, *o*. However, once a double heterozygote is created via outcrossing, a relevant recombination event can occur in any future generation via selfing as long as the genotype remains a double heterozygote. Because it takes, on average, two generations of selfing before heterozygosity at a site is lost, there are two generations in which meaningful recombination can occur following each outcrossing event (so we have 2*or* in [15] but *σr* in [5]). However, Denis Roze has pointed out that [14] is derived from [4], which assumes tight linkage (*r* << 1). He has found (pers. comm.) that the “2” is reduced for loosely linked loci, becoming “1” for unlinked loci. That is, the recominbational difference between selfing and partial asexuals will disappear for unlinked loci.

In addition to these segregation and recombination differences, a third difference arises because the effective strength of selection is greater with selfing than asexual reproduction (*s* vs. *hs*). Stronger selection reduces the frequency of deleterious alleles. As there are fewer deleterious backgrounds to avoid in tracing back a focal neutral site’s ancestry, background selection is made weaker. However, stronger selection also means there is less opportunity to recombine away from a deleterious allele before it is removed by selection, enhancing background selection. Overall, stronger selection reduces background selection if the rate of biparental reproduction (and the opportunity for recombinational escape) is high relative to selection.

As a consequence of the three differences between [15] and [5], background selection can be much stronger in partial sexuals than in selfers when the rate of biparental reproduction is low (Fig. 4). However, the reverse may be possible if the net effect of stronger selection in selfers is to increase background selection, which can happen when the rate of biparental reproduction is high relative to selection.

**Figure 4.**
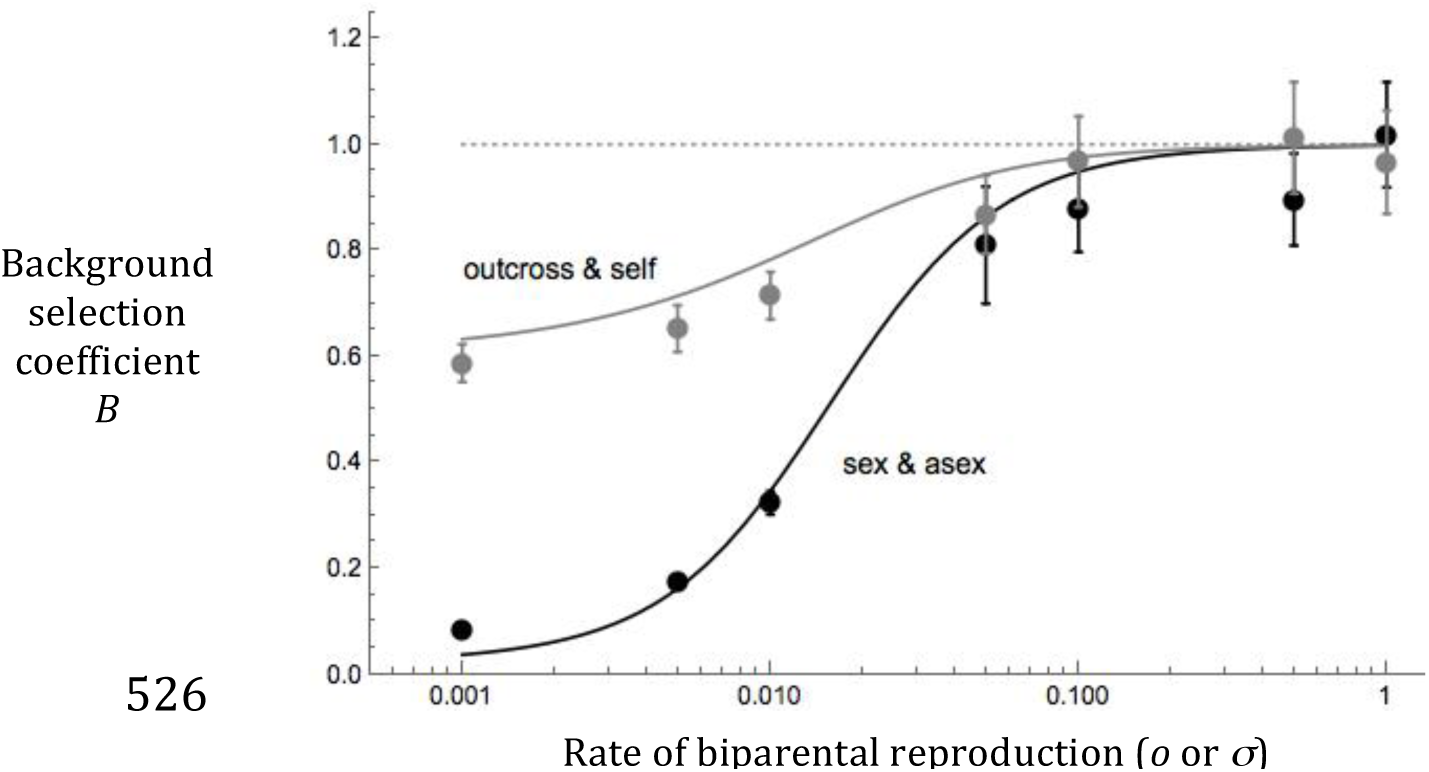
Background selection in systems with different forms of uniparental reproduction (asexuality and selfing). The background selection coefficient *B* is shown for systems with sexual and asexual reproduction (black) and systems with outcrossing and selfing (grey). Analytical approximations (lines) are contrasted with simulation results (points; error bars are 95% confidence intervals. The analytical approximation for selfing is based on an extension of [14] and is given in Kamran-Disfani & Agrawal (2014). Parameters: *U* = 0.02, *s* =0.02, *h* = 0.25, *L* = 10, (*N* = 10^4^ for simulations).

## Balancing Selection

With balancing selection, such as heterozygote advantage or negative frequency dependence, two alleles (*A*_1_ and *A*_2_) can be maintained indefinitely, i.e., balancing selection prevents normal coalescent processes from operating at the selected site and this affects coalescence at linked sites. Assuming the frequencies of the *A*_1_ and *A*_2_ (*p* and *q*, respectively) are stable, HUDSON and KAPLAN (1988) showed that closely linked neutral sites have an extended coalescent time under the assumption of obligate sexual reproduction. As mentioned in the previous section, the analysis becomes more complicated when the rate of sex is low because we must use an explicitly diploid perspective rather than classic haploid perspective. Here we cannot assume that the frequency of the *A*_2_/*A*_2_ genotype is negligible. As a result, there are 13 states in which we could find two alleles. In addition to the eight listed in the previous section, the additional five states are: (ix) both samples in a single *A*_2_*/A*_2_ individual; (x) both samples in separate *A*_2_*/A*_2_ individuals; (xi) one sample in an *A*_2_*/A*_2_ individual and the other on the *A*_1_-haplotype of an *A*_1_/*A*_2_ individual; (xii) one sample in an *A*_2_*/A*_2_ individual and the other on the *A*_2_-haplotype of an *A*_1_*/A*_2_ individual; and (xiii) one sample in an *A*_1_*/A*_1_ individual and the other sample in an *A*_2_*/A*_2_ individual.

In the previous section we ignored gene conversion for simplicity and because the persistence time of any given copy of the deleterious allele is short, limiting opportunity for gene conversion to be relevant. Here we include gene conversion because *A*_1_ and *A*_2_ are maintained indefinitely by selection so movement between genetic backgrounds via gene conversion may become important over long time scales, especially when the rate of sex is low so there is little chance for exchange via meiotic recombination. Specifically, we assume that (unbiased) mitotic gene conversion occurs at rate *γ*. This rate refers to gene conversion events that cover either the selected site or the neutral site but not both. (The model is built with an additional parameter for gene conversion events that cover both sites but such events do not affect the coalescent time.) Throughout, we use recombination to refer to (meiotic) cross-over recombination, in contrast to gene conversion.

In this analysis, we assume that the equilibrium frequencies of *A*_1_/*A*_1_, *A*_1_/*A*_2_, and *A*_2_/*A*_2_ genotypes are *P*_1/1_, *P*_1/2_, and *P*_2/2_. These frequencies are assumed to be stable and to change little in frequency within a generation (i.e., fitness differences at equilibrium are ignored). Without loss of generality, we use *P*_1/1_ = *p*^2^ + *C*_*A/A*_, *P*_1/2_ = 2(*pq – C*_*A/A*_), and *P*_2/2_ = *q*^2^ + *C*_*A/A*_ where *C*_*A/A*_ is the covariance in allelic state. Note *C*_*A/A*_ measures the same genetic property as *Fpq* but we use separate symbols because we use the relationship *F* = (1 – *o*)/(1 + *o*) for species without asexuality because the inbreeding coefficient is determined only by outcrossing rates, to a good approximation, provided selection is not too strong. With obligate sex (and no selfing) *C*_*A/A*_ = 0 but when there is partial asexuality, *C*_*A/A*_ can be either positive or negative.

Assuming *σ* and *r* are *O*(*ξ*) and *γ* and 1/*N* are *O*(*ξ*^2^), the expected coalescent time of two alleles sampled both from a single *A*_1_/*A*_1_ individual or from two separate *A*_1_/*A*_1_ individuals is found to be

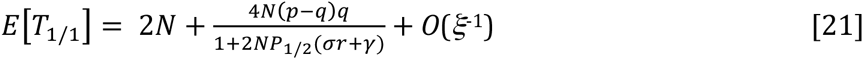

When both alleles are sampled from the *A*_2_ background, the coalescent time is the same as [21] but with *p* and *q* reversed. When two alleles are sampled from alternate backgrounds (*A*_1_ and *A*_2_), the expected coalescent time

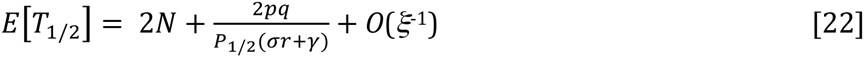

For the sake of discussion, we will focus on this latter result. In the case of obligate sex (*σ* = 1, *P*_1/2_ = 2*pq*), [22] simplifies to *E*[*T*_1/2_] = 2*N* + 1/(*r* + *γ*). The coalescent time is extended by the waiting time for the neutral site to move from one background to the other. This longer coalescent time should allow for the accumulation of more neutral variation surrounding sites under balancing selection (Hudson & Kaplan 1988). When the physical distance is small, *r* may be very small and this has led to the belief there may be a strong signature of balancing selection when gene conversion is ignored. However, for short physical distances, gene conversion may be much larger than recombination (*γ* >> *r*) and may play a more important role in coalescence (ANDOLFATTO and NORDBORG 1998; WIUF and HEIN 2000). As implied by [22], gene conversion weakens the signature of balancing selection and this is particularly true in partial asexuals (Fig. 6). At very short physical distances, mutation between *A*_1_ and *A*_2_ provides an important alternate route by which a neutral allele can switch sites, but this is not considered here. Our results apply to physical distances where *σr* + *γ* >> *μ*.

**Figure 6.**
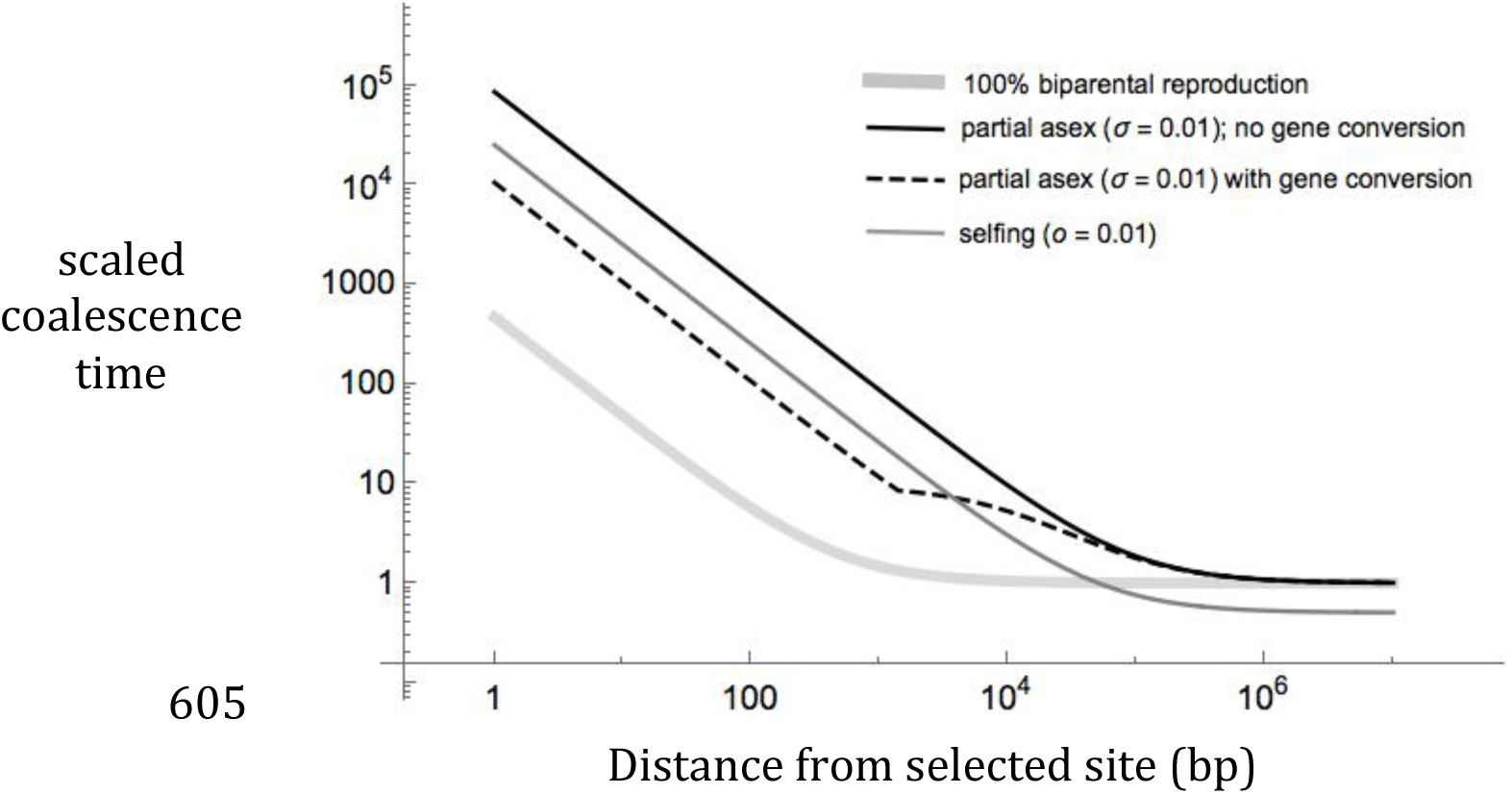
Coalescent times near a site under balancing selection. The expected time to coalescence (scaled to 2*N* generations) is shown as a function of the distance (in base pairs) from a site under balancing selection via heterozygote advantage (*W*_1/1_ *= W*_2/2_ *=* 1 – *s* and *W*_1/2_ = 1). The population is assumed to be fully sexual and outcrossing (eq. [22], *σ* = 1; thick gray line), mostly asexual (eq. [22], *σ* = 0.01; black line), or mostly selfing (eq. [24b], *o* = 0.01; thin gray line). We assume *r* = *ρ d* where *d* is the distance in base pairs from the selected site. For the partial asexual case, results are shown with and without gene conversion. For gene conversion, the model of Andolfatto and Nordborg (1998) is used: *γ* = *gd/L* for *d* < *L* and *γ* = *g* for *d* ≥ *L* where *L* is the length of gene conversion tracts. For the fully sexual case, the results with gene conversion are visually indistinguishable for those with *γ* = 0 for the parameter values used here. The selfing case assumes no gene conversion but gene conversion is not expected to affect coalescence in selfers (see Discussion). The selfing line asymptotes at 1/2 rather than 1 because of the direct effect of selfing on coalescence (Fig. 1). These results ignore mutation at the selected site and thus overestimate the coalescence time for extremely tightly linked sites (e.g., *d* << 100 if *μ* = 10^−9^, see text). Parameter values: *s* = 0.02, *N* = 10^5^, *ρ* = 10^−8^ (ASHBURNER 1989), *g* = 10^−6^ (SHARP & AGRAWAL, in review), *L* = 1500 (PRESTON & ENGELS 1996).

As expected, we find in [22] that partial asexuality reduces the effective rate of recombination from *r* to *σr*. However, there is an additional effect of asexuality that is not explicit in [22]; partial asexuality changes the equilibrium frequency of *A*_1_/*A*_2_ (i.e., *P*_1/2_ ≠ 2*pq*). The precise nature of this change depends on the mechanism of balancing selection. Consider balancing selection due to heterozygote advantage with symmetrical selection (i.e., *W*_1/2_ = 1, *W*_1/1_ = *W*_2/2_ = 1 – *s*). With obligate sex, at equilibrium *p* = *q* = ½ and *P*_1/2_ = ½. With partial asexuality and assuming the rate of sex is low, *p* = *q* = ½ but *P*_1/2_ = 2(*pq* – *C*_*A/A*_) where

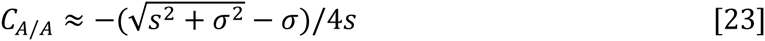

For example, with *s* = *σ* = 0.01, *P*_1/2_ ≈ 0.71. In this case, the frequency of heterozygotes (and thus the opportunity for recombination) is increased from 0.5 with full sex to 0.7 with partial sex. Even though *P*_1/2_ is higher, this does not cause an increase in the coalescence time because of the 99% reduction in meiotic recombination events when *σ* = 0.01. Equilibrium genotype frequencies under a more general model of balancing selection are provided in the supplementary material.

NORDBORG (1997) studied balancing selection in species with selfing. He found (ignoring gene conversion) that the expected coalescence time for sites on alternate backgrounds (*A*_1_ and *A*_2_) is

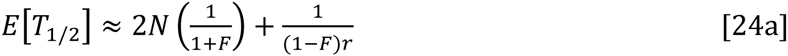

Assuming that rate of selfing is high (i.e., weak outcrossing, *o* << 1), this becomes

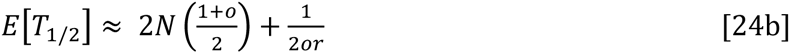

Again, the term in parenthesis is the direct effect of selfing. The second term is the increase in coalescent time due to linkage to the target of balancing selection. Contrasting this to the comparable term from the partial asexual result in [22], there are two differences. As noted in the background selection case, a little bit of outcrossing is twice as effective at allowing recombination as a little bit of sex (i.e., the effective rates of recombination are 2*or* vs. *σr*). Second, in partial asexuals the frequency of heterozygotes cannot be well approximated from the rate biparental reproduction and allele frequencies alone (as it is with selfing). Rather *C*_*A/A*_, which determines the frequency of heterozygotes through *P*_1/2_, is a function of both the rate of sex and the selection coefficients (as exemplified in [23]).

An important point missing from the comparison above is the likelihood of balancing selection. Consider the model of heterozygote advantage with *W*_1/2_ = 1, *W*_1/1_ = 1 – *s*, and *W*_2/2_ = 1 – *αs* where we assume *α* > 1 so that *W*_1/1_ > *W*_2/2_. Ignoring drift, polymorphism is always maintained in sex/asex systems. In contrast, with selfing, the maintenance of polymorphism requires that outcrossing not be too low, specifically that *o* > (*α* – 1)/(*α* + 1), otherwise fixation of the more fit homozygote *A*_1_/*A*_1_ is stable (see supplementary material for details). If one homozygote is more fit than another, variation will not be maintained by balancing selection in highly selfing species but will be in species with partial asexuality.

Of course, balancing selection can occur through mechanisms other than heterozygote advantage and in these other cases the contrast between partial asexuality and sex will be somewhat different. For example, consider a simple model of frequency dependent selection where the fitness of each genotype is negatively related to its frequency as follows *W*_1/1_ = 1 – (*p* – *α*)*s*, *W*_2/2_ = 1 + (*p* – *α*)*s*, and *W*_1/2_ = 1. Regardless of the reproductive system, at equilibrium *p* = *α* and the polymorphism is stable (i.e., in contrast to heterozygote advantage, there is no difference in the likelihood of balancing selection in this model between partial asexuality and selfing systems; see supplementary material). However with partial asexuality, there is no excess of heterozygotes (*C*_*A/A*_ ≈ 0), meaning that coalescence is only affected by the recombination effect (i.e., the reduction from *r* to *σr*) and there is no segregation effect. With other forms of negative frequency dependent selection (supplementary material), it is possible for there to be a deficit of heterozygotes with partial asexuality (*C*_*A/A*_ > 0), decreasing the opportunity for recombination beyond the change from *r* to *σr*, thereby further extending coalescent time.

## Discussion

We have examined how linked selection, either background or balancing selection, alters coalescence for pairs of alleles in partial asexuals. As expected, the lower level of genetic mixing in partial asexuals results in much stronger effects of linked selection. In diploids, partial asexuality affects coalescence both via segregation and recombination. The most obvious effect is through a reduction in the effective rate of recombination. In the case of background selection, the segregation effect can be understood as mediating how much a focal site is affected by deleterious alleles occurring on the homolog in addition to those on its own chromosome. If sex is low, it is equally affected by mutations on either chromosome, effectively doubling the mutation rate. In the case of balancing selection, the segregation effect is manifest in the way that partial asexuality affects the equilibrium frequency of heterozygotes at the selected locus, which regulates the opportunity for a neutral site to move between alternative selected haplotypes via recombination.

Ignoring linked selection, the direct effect of partial asexuality on coalescence is negligible unless the rate of sex is extremely small (i.e., on the order of 1/*N*; (BENGTSSON 2003; CEPLITIS 2003; HARTFIELD *et al.* 2015)). In contrast, there are strong effects of partial asexuality through background selection even when rates of sex are much larger than 1/*N*. Large reductions in *N*_*e*_ through background selection are expected under realistic mutation rates (*U* ≥ 0.1) if sex is below 10%.

When sex is high, only those mutations that are tightly physically linked make a substantial contribution to background selection. Across the genome of a highly sexual species, variation in gene density resulting in variation in the local deleterious mutation rate leads to variation in the strength of background selection, which contributes to variation in neutral diversity (CHARLESWORTH and CAMPOS 2014; ELYASHIV *et al.* 2014). These local differences may be detectable at the within-gene level (i.e., stronger background selection at the center of genes than on the edges, (LOEWE and CHARLESWORTH 2007; ZENG and CHARLESWORTH 2011)). When sex is low, deleterious alleles across the whole genome contribute to background selection. In fact, the majority of background selection experienced by a focal site is due to unlinked loci (Fig. 3B) but this is simply because unlinked sites are much more numerous. Even when sex is low, background selection is stronger from tightly linked sites than unlinked ones (Fig. 2B) so one might still expect to find differences in neutral diversity associated with gene density. However, the relationship between background selection and *r* is much weaker when sex is low, so gene density-diversity patterns are likely to be severely muted.

Because background selection can be so strong with uniparental inheritance, it may be more important than demographic history in determining within-population diversity. As has recently been discussed with respect to selfers (BRANDVAIN *et al.* 2013; BARRETT *et al.* 2014), transitions to uniparental inheritance can be associated with population bottlenecks but it can be difficult to infer whether reduced diversity in an evolutionarily young population is due to a small number of founders or the strong effects of background selection.

The analytical approximations not only quantify how much coalescence times are altered by the reproductive system, but they are also biologically interpretable. Hence these derivations provide insight into how segregation, recombination, and selection interact to affect average coalescence time. However, the approach used here cannot be used to make other, more detailed, predictions about *n*-sample genealogies. For example, although the major effect of background selection can be interpreted as a reduction in *N*_*e*_ (as done here), it has been shown that background selection also alters the shape of genealogies with relatively long external branches, leading to negative values of Tajima’s *D* (CHARLESWORTH *et al.* 1995; ZENG and CHARLESWORTH 2011; NICOLAISEN and DESAI 2012). Presumably, these effects would be magnified by low rates of sex.

The reduction in the effective rate of recombination with uniparental inheritance should make it much easier to detect balancing selection. Work by NORDBORG *et al.* (1996b) and NORDBORG (1997) showed that, in selfers, this is because (i) the physical distance over which the effects of balancing selection are realized is expanded and (ii) there is lower diversity within each selected allelic class because of stronger background selection. These arguments also apply to partial asexuals but there are differences between these alternative modes of uniparental reproduction. First, there is a higher effective rate of recombination with selfing than with partial asexuality for the same rate of uniparental reproduction (2*or* vs. *σr*, as discussed in the Results), which will make signatures of balancing selection weaker in selfers than partial asexuality (Fig. 6). There are also differences in the frequency of heterozygotes in partial asexuals compared to selfers but this likely plays a more minor role, conditional on the selected polymorphism being maintained. However, with heterozygote advantage, the conditions for a stable polymorphism are much broader in partial asexuals than selfers.

Another difference between partial asexuals and selfers is the importance of gene conversion. The frequency of heterozygotes puts an upper limit on the rate of exchange between allelic backgrounds via any genetic mechanism. Selfing directly reduces the frequency of heterozygotes, so gene conversion is not expected to be an important process (though it was not formally included in the model of (NORDBORG 1997)). In partial asexuals, heterozygotes can be very common even if sex is rare (especially if there is heterozygote advantage). Through asexual descent, a lineage can persist in heterozygous state for many generations, providing numerous opportunities for movement of linked sites between backgrounds via gene conversion. As implied by [22], gene conversion becomes the primary force erasing the signature of balancing selection when *σr* << *γ*, which must be true if sex is sufficiently rare. Because of this, the signature of balancing selection may be weaker in partial asexuals than selfers if the rate of biparental reproduction is very small (Fig. 6), even though the reverse is expected at less extreme levels of uniparental reproduction because of the difference in effective rates of recombination (2*or* vs. *σr*).

Here we explored how background and balancing selection affect coalescence times but we did not attempt to examine selective sweeps. In sexual populations, beneficial alleles spreading to fixation can drastically reduce coalesce times and diversity levels at closely linked sites (MAYNARD SMITH and HAIGH 1974; KAPLAN *et al.* 1989; STEPHAN *et al.* 1992; BARTON 1998; HERMISSON and PENNINGS 2005) and are believed to play an important role in shaping genomic patterns of diversity, at least in some species (ELYASHIV *et al.* 2014; WILLIAMSON *et al.* 2014). As with other forms of linked selection, we expect that the signature of selective sweeps will be more dramatic in systems with high rates of uniparental reproduction because of their reduced effective rates of recombination. For example, in the highly selfing *C. elegans*, sweeps are believed to have reduced variation across entire chromosomes (ANDERSEN *et al.* 2012). As with background and balancing selection, we speculate the “recombination effect” will not be the whole story in partial asexuals. With low rates of sexes, a “segregation effect” is likely because, under asexuality, the selected diploid genotypes are inherited intact (rather than their haploid gametes), altering the trajectory of the selected allele from that in the standard sexual (or haploid) model. The full consequences of partial asexuality with respect to selective sweeps remain a challenge for future work.

Though the consequences of sweeps are likely to be larger in systems with uniparental inheritance, sweeps are less likely to occur in such systems. Both theory (GLÉMIN and RONFORT 2012; KAMRAN-DISFANI and AGRAWAL 2014) and data (HOUGH *et al.* 2013; BURGARELLA *et al.* 2015) show that reductions in *N*_*e*_ due to background selection render selection less efficient, reducing the probability of fixation of beneficials, in selfers compared to outcrossers. A qualitatively similar effect is expected in partial asexuals.

A number of interesting and important diploid species reproduce predominantly asexually but these have been studied less by evolutionary geneticists than selfing species. Though there are similarities between partial asexuality and selfing, it is misleading to consider these two forms of uniparental inheritance as conceptually interchangeable. As population genomic data becomes easier to obtain, the hope is that such data will allow us to study the evolutionary consequences of different forms of uniparental reproduction and, ideally, gain insight into the evolutionary forces acting on reproductive mode. Understanding the coalescent properties of systems with uniparental inheritance is key to interpreting population genomic data and much work remains, especially with respect to partial asexuality.

## Acknowledgements

We thank Stephen I. Wright for extensive discussion and Denis Roze for sharing his unpublished result on background selection in selfers. This work was supported by the Natural Sciences and Research Council of Canada (AFA) and by a Marie Curie International Outgoing Fellowship, grant number MC-IOF-622936 project SEXSEL (MH).

